# Spectral power, but not phase connectivity, decodes the subanaesthetic ketamine state from surface EEG: the role of between-subject transferability

**DOI:** 10.64898/2026.07.21.739494

**Authors:** Hannes Schätzle, Frederic von Wegner

## Abstract

Subanaesthetic ketamine alters the content of consciousness while leaving responsiveness intact. We asked whether this state can be decoded from single eyes-closed EEG epochs, and how spectral power and phase-based connectivity compare when used as features. Re-analysing openly available 62-channel EEG from ten participants (awake versus subanaesthetic ketamine), we trained classifiers under leave-one-subject-out cross-validation to discriminate the two conditions from log band power, weighted phase-lag index (wPLI) connectivity, and their concatenation. Band power decoded the ketamine state above chance (balanced accuracy 0.71), whereas wPLI connectivity computed on the same epochs was at chance (0.47), and combining the feature sets did not improve on power alone. The dissociation held across three classifier families and across spatial montages, and was not explained by the dimensionality of the connectivity feature space. To identify its source, we decomposed the per-feature drug effect into components shared across subjects and subject-specific. The ketamine effect on connectivity was large within individuals but largely subject-specific, and therefore not transferable to held-out subjects (shared fraction 0.05), whereas the spectral effect was substantially shared across subjects (0.53); both feature classes carried comparable individual identity, so the asymmetry reflects transferability rather than fingerprint-likeness. The spectral signature was also recoverable from a sparse electrode subset, as a five-channel lateral montage performed as well as the full array. Undirected phase connectivity thus fails to decode subanaesthetic ketamine not through insufficient spatial sampling, but because the connectivity drug effect is subject-specific in form.

## 1 Introduction

Ketamine occupies an unusual position among psychoactive drugs. As a non-competitive *N* -methyl-D-aspartate (NMDA) receptor antagonist, it acts largely outside the GABAergic mechanisms of most general anaesthetics, and its effects are strongly dose-dependent: at anaesthetic doses it produces unconsciousness, whereas at subanaesthetic doses it induces a dissociative, psychedelic-like state while responsiveness and the general capacity for consciousness remain largely intact (Farnes et al 2020; Muthukumaraswamy et al 2015). This dissociation between altered conscious *content* and preserved conscious *level* makes subanaesthetic ketamine a valuable probe of how pharmacological perturbation reshapes ongoing brain dynamics, and has motivated a sustained search for electroencephalographic (EEG) signatures that track the ketamine state.

At the level of spectral power, the subanaesthetic ketamine signature is well characterised and highly reproducible across recording modalities: decreased posterior alpha power, increased frontal-midline theta power, and broadband increases in gamma power (Muthukumaraswamy et al 2015). Phase-based functional connectivity has been proposed as a complementary, and potentially more mechanistic, marker. The most consistently reported connectivity effect is a reduction in frontoparietal coupling; however, the clearest demonstrations of this effect, including reductions in the weighted and directed phase-lag index in the alpha band, have been obtained at *anaesthetic* doses, at or beyond the loss of consciousness (Vlisides et al 2017; Blain-Moraes et al 2014); in propofol sedation (a GABAergic agent, mechanistically distinct from ketamine), reductions in frontoparietal phase coupling track behavioural responsiveness itself (Chennu et al 2016). At subanaesthetic doses the connectivity picture is subtler and more often expressed as directional or effective rather than undirected coupling (Muthukumaraswamy et al 2015), leaving open the question of how much undirected phase-based connectivity the conscious, subanaesthetic state actually carries.

The weighted phase-lag index (wPLI) is a well-established measure for functional connectivity because it quantifies consistent non-zero phase lags between signals while being insensitive to the zero-lag synchrony introduced by volume conduction and common references (Vinck et al 2011). Connectivity-based representations have been shown to discriminate the ketamine state from control in other modalities and analysis frameworks, for example, whole-brain functional connectivity classifies ketamine from saline with high accuracy in pharmacological fMRI (Joules et al 2015), and connectivity- and graph-theoretic features support cross-state classification that includes subanaesthetic ketamine (Jang et al 2024). Within scalp EEG, however, the evidence is more equivocal. In the closest comparable design, low-density EEG during subanaesthetic ketamine, conventional pairwise connectivity was a comparatively weak discriminator, and a robust, classifiable signal emerged only once higher-order, multivariate interactions were considered (Herzog et al 2024).

Two features of the prediction problem complicate the move from group-level connectivity differences to single-trial decoding. First, most connectivity evidence is established through group-level contrasts, which need not imply that the same features support prediction in held-out individuals. A difference that is reliable on average can fail to generalise if it is not consistent in form across people. Second, this concern is especially acute for functional connectivity, whose structure is strongly subject-specific (Finn et al 2015; Gratton et al 2018). In functional MRI, individual connectivity profiles are stable enough to act as a “fingerprint”, with the frontoparietal network, the network most implicated in ketamine’s effects, among the most individuating, and the connections that discriminate individuals largely distinct from those that track states (Mantwill et al 2022). Under a subject-aware evaluation, in which no participant contributes to both training and test data, a connectivity-based classifier can therefore succeed only to the extent that the drug effect is shared across individuals rather than subject-specific.

Here we re-analyse the high-density (62-channel) EEG dataset of Farnes et al, in which ten healthy volunteers were recorded at rest before and during a continuous subanaesthetic ketamine infusion (Farnes et al 2020). Whereas the original study characterised the ketamine state using measures of signal diversity, we ask a distinct question: whether the ketamine state can be classified at the level of individual eyes-closed epochs, and how single-channel spectral and pairwise phase-based connectivity features compare when used for this purpose.

## 2 Materials and Methods

### 2.1 Dataset and participants

We re-analysed a publicly available scalp-EEG dataset acquired by Farnes and colleagues (Farnes et al 2020), and shared via the Dryad repository (DOI: 10.5061/dryad.j9kd51c9q). The dataset comprises high-density EEG recordings from *N* = 10 healthy adult volunteers (7 male, 3 female participants; median age 27.5 years, range 21–44) acquired in an open-label, within-subject design comparing normal wakefulness (awake; AW) to a subanaesthetic ketamine-induced psychedelic state (KET). The original study was approved by the Norwegian Regional Committees for Medical and Health Research Ethics (reference 2015/1520/REK Sør-Øst A); written informed consent was obtained from all participants by the original investigators. The present study used only de-identified, pre-processed recordings released by the original authors and required no additional ethical approval.

Recordings were collected on a single experimental day at the Intervention Centre of Oslo University Hospital, comprising four spontaneous EEG runs acquired in a fixed order: (i) awake, eyes open; (ii) awake, eyes closed; (iii) ketamine, eyes open; (iv) ketamine, eyes closed. Each spontaneous run lasted approximately two minutes. Racemic ketamine (Ketalar^®^, Pfizer, Norway; 10 mg mL^−1^) was administered as a continuous intravenous infusion using a syringe pump (B. Braun Perfusor Space), ramped from 0.1 to a maximum of 1.0 mg kg^−1^ h^−1^ in 0.1 mg kg^−1^ h^−1^ steps every five minutes, then stabilised once both participant and attending anaesthesiologist agreed that a noticeable subjective effect had been reached. Across participants, the median maintenance infusion rate was 0.7 mg kg^−1^ h^−1^ (range 0.5–1.0) and the median total infusion time was 43.5 minutes (range 37–73); these doses produced reliable dissociative effects without loss of responsiveness or consciousness. The present analyses use only the two eyes-closed runs per participant, so that group-level contrasts are not confounded by the well-known effects of eye opening on posterior alpha rhythms and other components of the spontaneous EEG; each participant thus contributes one awake-EC and one ketamine-EC recording, for a total of 20 recordings analysed.

### 2.2 EEG acquisition

Continuous EEG was acquired with two TMS-compatible 32-channel BrainAmp DC amplifiers (Brain Products GmbH, Germany) connected to a TMS-compatible EEG cap arranged according to the international 10–10 system, with a forehead common reference and a scalp ground. Electrode impedances were maintained below 10 kΩ. Signals were digitised at 5000 Hz with 16 bit resolution and a 1000 Hz hardware antialiasing low-pass filter. The shared recordings contain 62 scalp channels (all labelled as EEG in the standard 10–10 montage, ranging from Fp1/Fpz/Fp2 anteriorly to O1/Oz/O2 and Iz posteriorly); no electro-oculogram or auxiliary channels are present in the released files. All 62 channels were retained for analysis.

### 2.3 Preprocessing

The Dryad-released recordings had already been preprocessed by the original authors using EEGLAB. Their pipeline, summarised here for completeness and described in full in Farnes et al (2020), comprised: (i) segmentation of the continuous recording into non-overlapping 8 s epochs (15 epochs per recording); (ii) marking and interpolation of channels showing flat, noisy or abnormal-amplitude traces; (iii) rejection of epochs with high variance, large transient deflections or movement artefacts; (iv) zero-mean baseline correction across the full epoch; (v) re-referencing to the common average; (vi) zero-phase Hamming-window FIR band-pass filtering between 0.5 Hz and 45 Hz (default filter order 3 × rate*/*2); (vii) downsampling to 250 Hz; (viii) manual rejection of independent components corresponding to ocular artefacts; and (ix) computation of the surface Laplacian, which subtracts each electrode’s mean-neighbour signal to yield an estimate of the local current source density (CSD), thereby attenuating volume-conduction effects inherent to scalp EEG (Hjorth 1975).

After verifying that no epoch in the released data exceeded the 250 µV peak-to-peak threshold, all 276 epochs across the 20 eyes-closed recordings were retained for analysis (median 14 epochs per recording, range 11–17; per-recording counts in Table 5). All downstream analyses were performed in Python 3.11 with MNE-Python v1.7 (Gramfort et al 2013).

### 2.4 Spectral analysis

For descriptive comparison of the two states, we computed the power spectral density (PSD) of each epoch using Welch’s method (mne.compute psd, method = “welch”) with a 2 s Hann window, 50% overlap and FFT length matched to the window. The analysed range was 1 Hz to 40 Hz. Per-recording PSDs were obtained by averaging across epochs, then across channels, yielding one spectrum per subject and condition. Group-level grand-average spectra are reported as the mean across subjects with ± 1 SEM, with SEM computed across subjects (the unit of replication). To visualise condition-specific changes independently of inter-subject variation in absolute power, we additionally computed a per-subject log power ratio log *P*_ket_(*f*) − log *P*_aw_(*f*) and averaged this across subjects. Canonical frequency bands were defined as delta (1–4 Hz), theta (4–8 Hz), alpha (8–13 Hz) and beta (13–30 Hz). For the per-band comparison, power within each canonical band was averaged across all 62 electrodes and all retained epochs per subject and condition, then log-transformed, yielding one value per subject, condition, and band.

### 2.5 Feature extraction for classification

Three feature sets were constructed from eyes-closed epochs and used as input to the decoding analysis. The unit of observation throughout was the individual 8-second epoch.

#### Set A: spectral power

For each epoch, a Welch PSD was estimated with the FFT length set to the full epoch (2000 samples), with no overlap. Mean power in each of the four canonical bands was then averaged across all channels (global) and within each of five anatomical regions assigned from electrode names (frontal: Fp*, AF*, F*; central: C*; parietal: P*; occipital: O*; temporal: T*). All resulting powers were log-transformed, yielding 24 features per epoch.

#### Set B: phase-based functional connectivity

For each epoch, weighted phase-lag index (wPLI; Vinck et al (2011)) was estimated between all 62 × 62 electrode pairs in three frequency bands (theta, alpha, beta). Delta (1–4 Hz) was omitted from the connectivity analysis because the 1 s sub-windows used for wPLI estimation (see below) span only one to four delta cycles, too few to support a stable phase-lag estimate at these frequencies. wPLI is a phase-based connectivity measure designed to be insensitive to zero-lag synchrony arising from common-source artefacts and from volume conduction. Because the underlying signals here are already surface-Laplacian-transformed (Section 2.3), the resulting wPLI values quantify phase coupling between local CSD estimates rather than between scalp potentials, and are expected to be still less contaminated by residual volume-conduction effects than scalp-level wPLI would be. Because a stable wPLI estimate requires multiple trials, each 8 s epoch was internally re-segmented into 1 s sliding sub-windows with 0.5 s step (≈ 15 sub-windows per epoch); these sub-windows were then passed as the trial dimension to `mne_connectivity.spectral_connectivity_epochs` with `method = “wpli”` and a multitaper spectral estimator (`mode = “multitaper”, faverage = True`). The resulting 62 × 62 matrix was symmetrised, the diagonal set to zero, and the upper triangle vectorised, giving 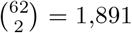 edges per band and 5,673 features per epoch in total.

#### Set C: combined

Set C is the epoch-level concatenation of Sets A and B (24 + 5,673 = 5,697 features), included to test whether spectral power and inter-channel phase coupling carry complementary information about the ketamine state.

### 2.6 Channel subsets

To assess how much the electrode montage can be reduced without losing the discriminative signal, the feature pipelines above were repeated on four anatomically motivated subsets of the montage that approximate clinically deployable configurations:

- **Full 62-channel** montage (baseline; results reused from the main analysis).
- **Both-lateral** (10 channels: Fp1, F7, T7, P7, O1 and Fp2, F8, T8, P8, O2), the electrode positions making up the two longitudinal temporal chains of the standard clinical longitudinal bipolar (“banana”) montage. Note that we use these electrode *locations* only; all features are computed on the surface-Laplacian/CSD estimates described in Section 2.3, not on bipolar derivations.
- **Left-lateral** (5 channels: Fp1, F7, T7, P7, O1) and **Right-lateral** (5 channels: Fp2, F8, T8, P8, O2).
- **BIS-Quatro** (4 channels: Fpz, Fp1, AF3, F7), an approximation of the four-electrode forehead strip of the Bispectral Index (BIS, Medtronic/Covidien) monitor used in routine anaesthesia practice; we placed the four virtual sensors at the nearest available 10-10 positions to the BIS Quatro ES_1_–ES_4_ pads.

For each subset, Set A was rebuilt as per-channel log band power in the four canonical bands (4 × *n*_ch_ features), and Set B was extracted from the previously-computed full-montage epoch-level wPLI matrices, sub-indexed to the channels of the subset 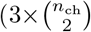 features). Sub-indexing already-computed matrices, rather than recomputing wPLI on the reduced montage, ensures that the underlying connectivity estimates are identical across subsets so that performance differences are attributable to channel selection alone.

### 2.7 Classification pipeline

Three classifier families were trained to discriminate ketamine (KET) from awake (AW) epochs, separately for each feature set: random forests (RF; (Breiman 2001)), support vector machines with a radial basis function kernel (SVM)(Chang and Lin 2011), and gradient-boosted trees (XGBoost)(Chen and Guestrin 2016). The random forest is the primary classifier; the SVM and XGBoost serve as robustness checks, establishing that the spectral–connectivity dissociation reflects the features rather than the choice of classifier. At the epoch level the two classes were close to balanced (135 ketamine, 141 awake). To prevent within-subject leakage, all cross-validation grouping used subject id, so that no subject appeared simultaneously in training and test data.

#### Cross-validation

All models were evaluated with leave-one-subject-out (LOSO) outer cross-validation. Each of the ten subjects served in turn as the held-out test set while the classifier was trained on the remaining nine. Hyperparameters were tuned within each outer training set by an inner four-fold GroupKFold, using GridSearchCV scored by balanced accuracy. The inner split received only the training fold’s subject labels, so the held-out subject was never touched during tuning; we verified zero subject overlap between inner-training and inner-validation folds.

#### Full 62-channel models

For the wPLI feature sets (Set B, 5,673 features; Set C, 5,697 features), which greatly exceed the 276 epochs, a shared PCA step (`sklearn.decomposition.PCA`, 100 components) preceded the classifier, fitted strictly inside the inner CV folds within an `sklearn.pipeline.Pipeline`, so that no information from the test fold influenced the PCA basis and all three estimators operated on identical inputs. Set A used no PCA. The SVM pipeline additionally included a `StandardScaler`, likewise fitted inside the inner folds. Estimator-specific settings were: RF with `class_weight = “balanced_subsample”`; SVM with `class_weight = “balanced”` and ROC-AUC computed from the decision function; XGBoost with `tree_method = “hist”` and no class reweighting. Hyperparameter grids were: RF (720 combinations) over *n*_estimators_ ∈ {200, 400, 600, 1000}, max depth ∈ {5, 10, 20}, min samples split ∈ {2, 4, 8, 10}, min samples leaf ∈ {1, 2, 3, 4, 5}, and max features 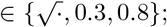 SVM (12 combinations) over *C* ∈ {0.1, 1, 10, 100} and *γ* ∈ {scale, 0.01, 0.001}; XGBoost (∈ {32} combinations) over *n*_estimators_ ∈ {300, 600}, max depth ∈ {3, 6}, learning rate ∈ {0.05, 0.1}, subsample ∈ {0.8, 1.0}, and colsample bytree ∈ {0.8, 1.0}. All estimators used a fixed random seed.

#### Channel-subset models

Because the channel subsets reduce feature dimensionality by one to two orders of magnitude, no PCA was applied. Grids were smaller: RF (243 combinations) over *n*_estimators_ ∈ {200, 400, 800}, max_depth 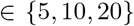, min_samples_split ∈ {2, 6, 10}, min_samples_leaf ∈ {2, 4, 5}, and max_features 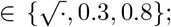; SVM as for the full montage; XGBoost (8 combinations) over *n*_estimators_ ∈ {200, 400}, max_depth ∈ {3, 6}, and learning_rate ∈ {0.05, 0.1} (subsample and colsample bytree fixed at 1.0). The outer LOSO scheme was identical to the full-montage analysis.

#### Evaluation metrics

For each outer fold we report balanced accuracy and ROC-AUC, averaged across the ten LOSO folds. Balanced accuracy is the primary metric because the two classes are not perfectly balanced at the epoch level. Because each LOSO fold comprises a single held-out participant, across-fold variance is inherently high; significance is therefore established by the permutation test on the across-fold mean (Section 2.8), not by the fold-to-fold spread.

### 2.8 Statistical evaluation

Per-band differences in log power between conditions were tested separately with two-sided paired Wilcoxon signed-rank tests across the ten subjects, with *p*-values Holm-Bonferroni-corrected across the four bands.

To test whether each (subset × feature-set × classifier) cell performed significantly above chance, we constructed a label-shuffled null distribution for each model. Within each permutation, recording-level condition labels were shuffled *within subjects* (so that each subject retained one awake and one ketamine recording, but with their assignment randomised); all epochs inherited the new recording-level label. This preserves the subject grouping required by the leave-one-subject-out scheme and the within-subject paired structure, but breaks the association between the EEG features and the condition labels. For each permutation, the pipeline was refit on each outer fold using the hyperparameters previously selected on the unpermuted data (so that hyperparameter search was not repeated for every permutation), and the balanced accuracy and ROC-AUC averaged across folds were recorded. The procedure was repeated for *N*_perm_ = 1,000 permutations. Two-sided permutation *p*-values were computed as

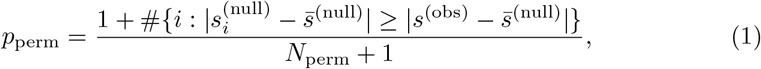

where *s* denotes the metric of interest (balanced accuracy or ROC-AUC) and 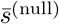 is the mean of the null distribution. With *N*_perm_ = 1,000, the minimum reportable *p*-value is 1*/*1001 ≈ 0.001. No multiple-comparison correction was applied across cells, which are highly non-independent (sharing epochs and feature extractors); individual *p*-values are reported for each cell.

### 2.9 Feature importance and back-projection to edge space

To interpret which inter-channel interactions drove the wPLI-based classifier (Set B on the full montage), we projected the random-forest feature importances from the 100-dimensional PCA space back into the original 5,673-edge space. Importances were the standard mean decrease in impurity (Gini importance), averaged over all trees in the forest (Breiman 2001). Node impurity quantifies how mixed the class labels are among the samples reaching a given node: it is zero when a node contains only one class and maximal when the classes are equally represented, and is measured here by the Gini index, 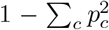, where *p*_*c*_ is the proportion of class *c* at the node. Each split is chosen to reduce this impurity, and the importance of a feature is the total impurity reduction it produces, summed over all splits that use it and averaged across the forest (Louppe et al 2013). For each outer fold *f* we obtained the matrix of PCA loadings *V* ^(*f*)^ ∈ ℝ^100*×*5,673^ and the vector of RF feature importances *I*^(*f*)^ ∈ ℝ^100^, and computed the back-projected importance for edge feature *j* as

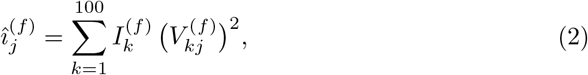

i.e., a weighted sum of squared loadings; squaring removes the sign ambiguity that affects principal-component loadings across folds. The per-edge importance reported in Figure 8 is the mean of 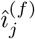 across the ten outer folds. Band-specific importance maps were obtained by restricting the sum to the 1,891 edges belonging to each frequency band; node-level importance was computed as the sum of incident edges. We additionally report the top-20 ranked edges by mean importance, for descriptive interpretation only; these are not statistically tested.

### 2.10 Variance decomposition and drug-effect transferability

To characterise how the ketamine effect on each feature class is distributed across and within subjects, we decomposed, for every feature, the per-subject drug effect into a component shared across subjects and a subject-specific component, separately for the 5,673 wPLI edges and the 24 band power features. This follows the logic of variance-of-treatment-effect decompositions (Ding et al 2019) and of intraclass-correlation reliability coefficients (Shrout and Fleiss 1979). The specific transferability ratio defined below (Eq. 5) is, to our knowledge, not a standard named coefficient; we construct it here from these two established ideas and apply it per feature to the awake-versus-ketamine contrast. We use “transferability” here in a statistical sense (the cross-subject-shared fraction of an effect), not in the sense of transfer learning in machine learning.

For feature *j* and subject *s* (*s* = 1, …, *S* with *S* = 10), let 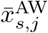 and 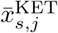 denote the mean feature value over that subject’s awake and ketamine epochs, respectively, and let

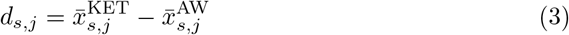

be the subject-specific drug effect. We summarise the across-subject distribution of *d*_*s,j*_ by its mean and variance,

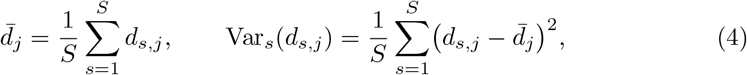

where 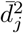 is the *shared* (cross-subject-consistent) component of the drug effect and Var_*s*_(*d*_*s,j*_) is the *subject-specific* component. We then defined the drug-effect *transferability* of feature *j* as the proportion of the total per-subject drug-effect energy that is common across subjects,

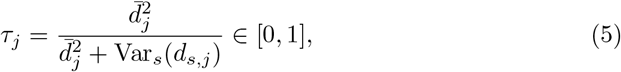

where the numerator 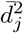 measures the squared *shared* effect and the denominator adds the across-subject variance of the per-subject effects. Thus *τ*_*j*_ = 1 when every subject shows an identical drug effect (no subject-specific component), and *τ*_*j*_ = 0 when the per-subject effects average to zero so that the entire effect is subject-specific. Because a classifier evaluated on held-out subjects can exploit only the shared component, *τ*_*j*_ quantifies how much of a feature’s drug effect is available for cross-subject generalisation. By construction it is an intraclass-correlation-like ratio of a shared component to a shared-plus-subject-specific total, computed on the per-subject effects rather than on raw scores.

We additionally computed, per feature: the between-subject variance 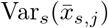 of the per-subject condition means, as a measure of how much the feature varies with subject identity; the subject intraclass correlation ICC_subject_ (consistency form, ICC(3,1); Shrout and Fleiss (1979)), the proportion of total variance attributable to subject identity; and the ratio of between-subject variance to the mean squared per-subject drug effect 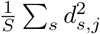, which falls below one wherever the within-subject drug effect exceeds the between-subject variance. Together with *τ*, these quantities separate the size of the between-subject (identity) variance from the cross-subject consistency of the drug effect.

The band power decomposition used the identical procedure as the wPLI decomposition, verified to reproduce the wPLI values when applied to the wPLI features; band power–wPLI comparisons of *τ* are band-matched to the theta/alpha/beta range covered by wPLI (Section 2.5).

To verify that connectivity carried individual identity, we trained a random-forest classifier to predict subject identity (*S* = 10 classes) from each feature set, evaluated across drug conditions (train on awake epochs, test on ketamine epochs, and vice versa) so that identity decoding could not exploit within-recording structure. Balanced accuracy was compared against the 1*/S* = 0.10 chance level.

### 2.11 Software and code availability

Analyses were implemented in Python 3.11 using NumPy 1.26, SciPy 1.13, pandas 2.2, scikit-learn 1.5, MNE-Python 1.7 (Gramfort et al 2013) and MNE-Connectivity 0.7. All code is available on https://github.com/05d762de69/ketamine-eeg-prediction organised as numbered Jupyter notebooks corresponding to each analysis step.

## 3 Results

### 3.1 Resting EEG power in the awake and ketamine conditions

Grand-average power spectral densities, computed across all 62 electrodes and the ten participants, are shown in Figure 1. Both conditions showed the characteristic eyes-closed morphology, which is a low-frequency 1*/f* -like roll-off with an alpha peak near 10 Hz, followed by a shallower decline through the beta range.

The within-subject log power ratio log *P*_ket_(*f*) − log *P*_aw_(*f*) differed significantly from zero in two contiguous frequency clusters, in both of which power was lower under ketamine: a low-frequency cluster spanning 1.5–9 Hz (*p* = 0.014) and a beta-range cluster spanning 13.5–24.5 Hz (*p* = 0.004). Significant clusters are marked in Figure 1.

**Fig. 1.**
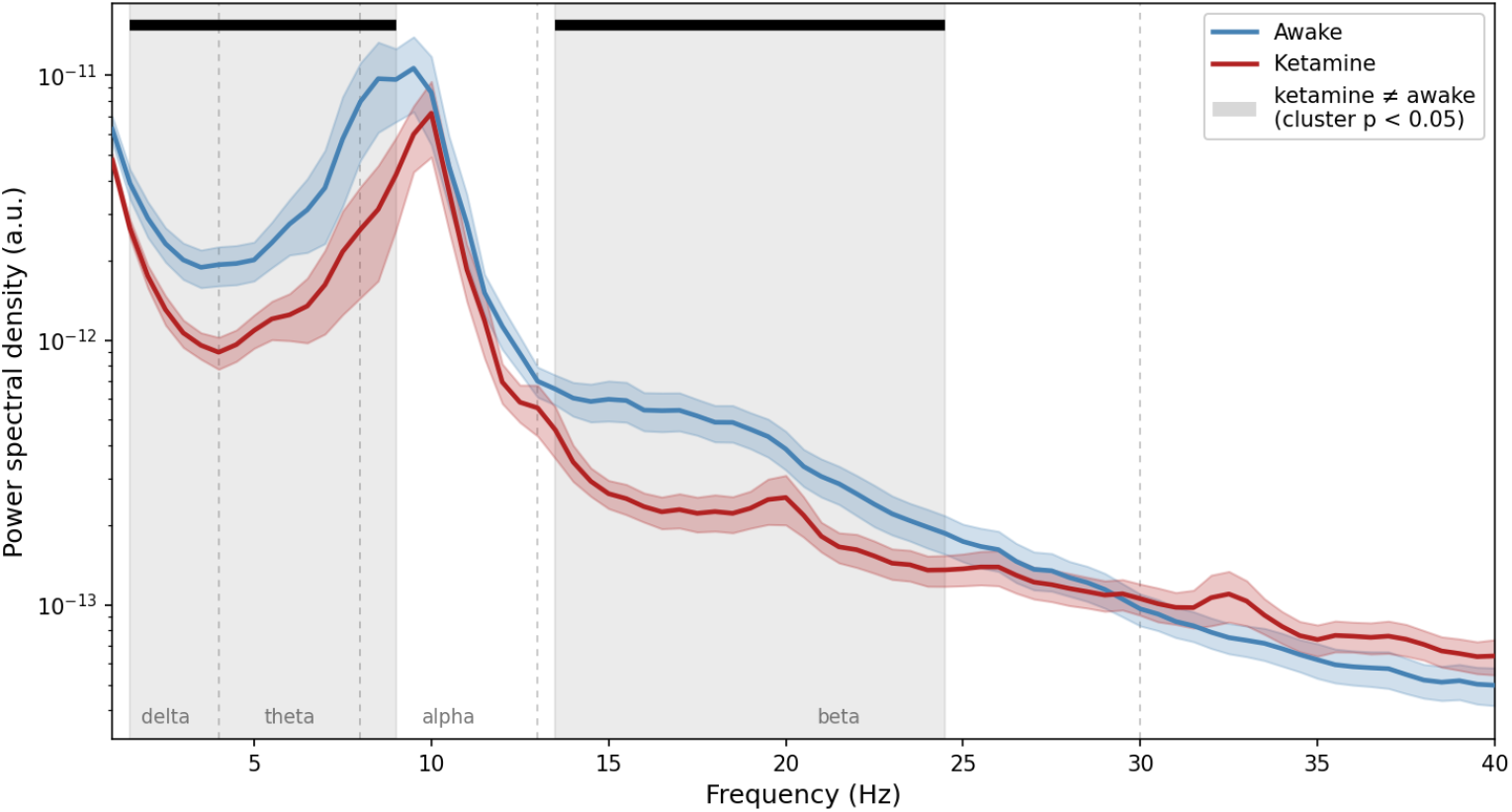
Grand-average power spectral density for the awake (blue) and ketamine (red) eyes-closed conditions. Solid lines, across-subject mean; shaded bands, *±* 1 SEM. Dashed vertical lines mark the canonical frequency-band boundaries (labelled below the axis); grey shading and the black horizontal bars mark the frequency ranges of significant between-condition clusters. Power is in arbitrary units (current-source-density estimates), so only relative differences are interpretable

Per-band log power differences are reported in Table 1. Power was lower under ketamine in all four bands after Holm correction: theta (mean log difference −0.71, a ∼50% reduction in linear power, *p*_Holm_ = 0.008), beta (−0.48, ∼38%, *p*_Holm_ = 0.008), alpha (− 0.64, ∼ 47%, *p*_Holm_ = 0.039), and delta (− 0.37, ∼ 31%, *p*_Holm_ = 0.039). All ten subjects decreased in theta and beta, nine of ten in alpha, and eight of ten in delta (Table 1).

**Table 1.** Per-band paired Wilcoxon signed-rank tests on the within-subject log power difference (ketamine − awake). *n*_+_*/n*_−_: number of subjects with a positive / negative log difference; *p*_Holm_: Holm-Bonferroni-corrected across the four bands.

| Band | Range (Hz) | Mean $\Delta \log$ | Median $\Delta \log$ | $n_+/n_-$ | $W$ | $p$ | $p_{\text{Holm}}$ |
| --- | --- | --- | --- | --- | --- | --- | --- |
| Delta | 1–4 | $-0.37$ | $-0.43$ | 2/8 | 5.0 | 0.020 | 0.039 |
| Theta | 4–8 | $-0.71$ | $-0.70$ | 0/10 | 0.0 | 0.002 | 0.008 |
| Alpha | 8–13 | $-0.64$ | $-0.81$ | 1/9 | 7.0 | 0.037 | 0.039 |
| Beta | 13–30 | $-0.48$ | $-0.42$ | 0/10 | 0.0 | 0.002 | 0.008 |

Per-subject spectra (Figure 4) showed the reduction was present in every participant.

### 3.2 Classification performance on the full 62-channel montage

#### Full-montage band power (Set A)

With the random forest, Set A achieved a mean balanced accuracy of 0.707 (SD 0.191) and a mean ROC-AUC of 0.803 (SD 0.269) across the ten LOSO folds, exceeding chance on two-sided permutation tests on both metrics (*p* = 0.011 and *p* = 0.010). Averaged across folds, sensitivity was 0.646 and specificity 0.768 (Table 2). Performance varied markedly across held-out participants: Two subjects fell near or below chance while the remainder ranged up to 1.00 (Figure 5), examined further in Section 3.3.

#### Full-montage wPLI (Set B)

The 5,673-edge wPLI feature vector, reduced to 100 PCA components fitted within the inner CV folds, yielded a mean balanced accuracy of 0.474 (SD 0.165) and a mean ROC-AUC of 0.489 (SD 0.202), neither distinguishable from chance (*p* = 0.563 and *p* = 0.853). Averaged across folds, sensitivity was 0.475 and specificity 0.473 (Table 2).

#### Full-montage combined features (Set C)

Set C reached a mean balanced accuracy of 0.671 (SD 0.097) and ROC-AUC of 0.820 (SD 0.068), above chance (*p* = 0.012 and *p* = 0.002). Its balanced accuracy fell below that of Set A, while its ROC-AUC was comparable (Table 2).

**Table 2.**
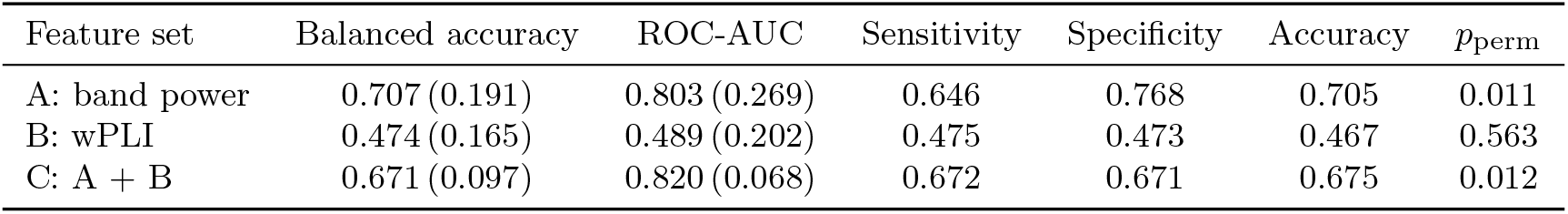
Full-montage classification performance for the primary random-forest classifier. Balanced accuracy and ROC-AUC are the mean across the ten leave-one-subject-out folds (SD in parentheses). Sensitivity and specificity are the mean across folds; accuracy is on the pooled predictions. *p*_perm_ values are two-sided permutation tests on balanced accuracy.

#### The dissociation holds across classifier families

The same pattern held for the SVM and XGBoost classifiers (Table 3). band power exceeded chance for all three (*p*_bacc_ = 0.007–0.012), wPLI was indistinguishable from chance for all three (*p*_bacc_ = 0.449–0.628), and the combined set did not exceed band power on balanced accuracy.

**Table 3.**
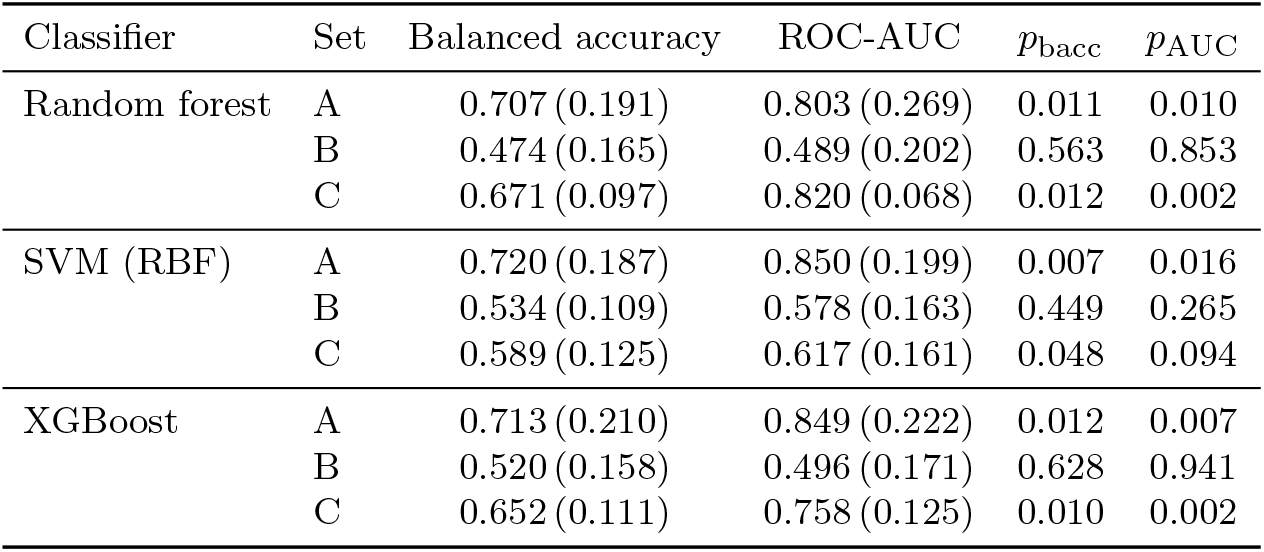
Full-montage classification performance across three classifier families. Balanced accuracy and ROC-AUC are the mean across the ten leave-one-subject-out folds (SD in parentheses); *p*_bacc_ and *p*_AUC_ are two-sided permutation *p*-values. Band power (Set A) exceeds chance and wPLI (Set B) is at chance for every classifier.

### 3.3 Classification performance across channel subsets

We repeated the classification analysis on the four channel subsets (Section 2.6) for the primary random-forest classifier. Figure 2 shows the mean balanced accuracy and ROC-AUC across the ten leave-one-subject-out folds for each (subset × feature set) cell, with two-sided permutation *p*-values; fold-level estimates are in Table 4.

**Fig. 2.**
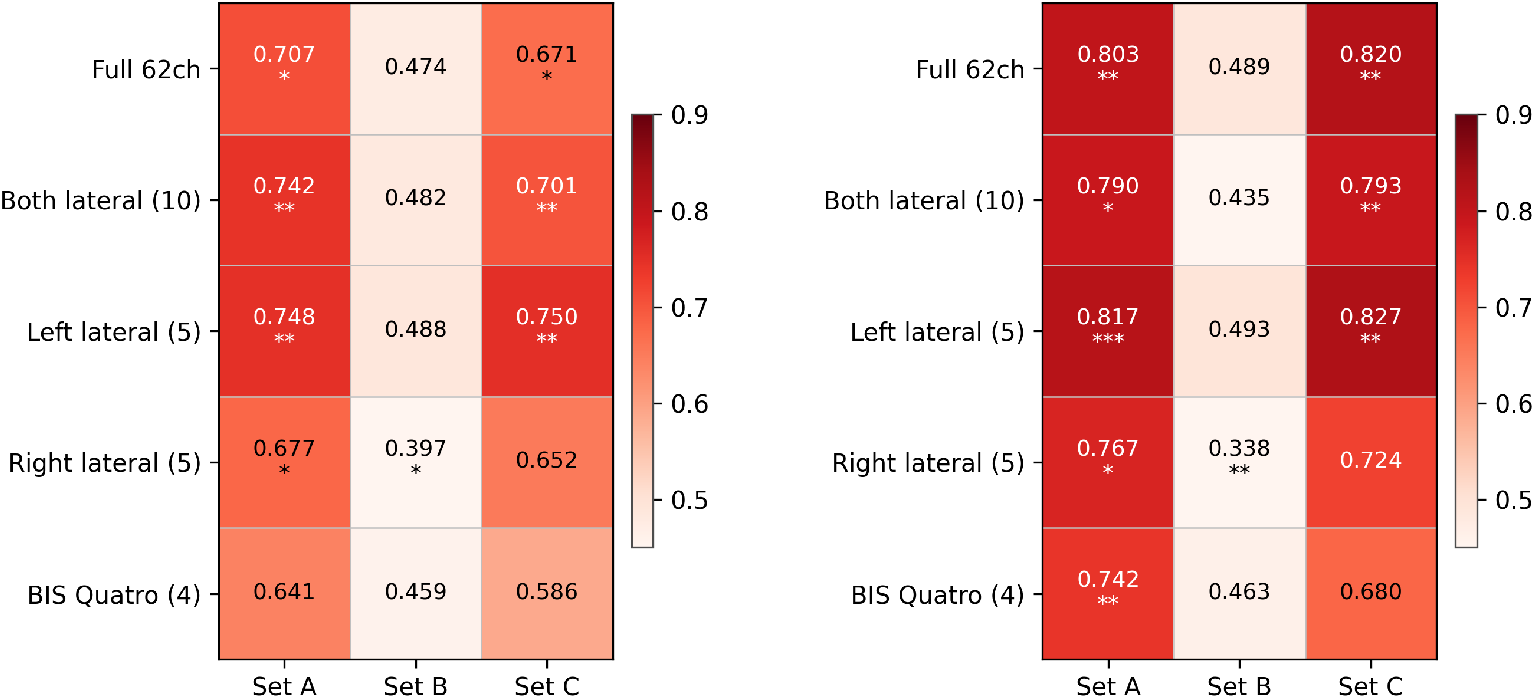
Classification performance across channel subsets and feature sets, for the primary random-forest classifier. Cells show the mean of balanced accuracy (left panel) and ROC-AUC (right panel) across the ten leave-one-subject-out folds. The Full 62-channel row reproduces the results of Section 3.2; remaining rows correspond to the channel-subset analysis. Within-cell symbols denote two-sided permutation *p*-values: ^*^ *p <* 0.05, ^**^ *p <* 0.01, ^***^ *p <* 0.001

#### Channel-subset performance, Set A (band power)

On Set A, the both-lateral 10-channel montage reached a mean balanced accuracy of 0.742 and ROC-AUC of 0.790; left-lateral reached 0.748 / 0.817 and right lateral 0.677 / 0.767 (balanced accuracy / ROC-AUC). All three lateral × Set A cells exceeded chance on balanced accuracy (both-lateral *p* = 0.002, left-lateral *p* = 0.007, right-lateral *p* = 0.017; Table 4).

#### Channel-subset performance, BIS Quatro (Set A)

The BIS Quatro montage yielded a mean balanced accuracy of 0.641 and ROC-AUC of 0.742 on Set A. ROC-AUC exceeded chance (*p* = 0.003); balanced accuracy did not (*p* = 0.056; Table 4).

#### Channel-subset performance, Set B (wPLI)

On Set B, balanced accuracy was at or below chance for every subset: both-lateral 0.482 (*p* = 0.701), left-lateral 0.488 (*p* = 0.818), and BIS Quatro 0.459 (*p* = 0.330). The right-lateral cell was again below chance (0.397, *p* = 0.031; Table 4). Back-projected edge importances for the full-montage Set B classifier are reported in Figure 8 and Table 7; as that classifier did not exceed chance, they are not interpreted further.

#### Channel-subset performance, Set C (combined)

Balanced accuracy for Set A versus Set C was 0.742 versus 0.701 (both-lateral), 0.748 versus 0.750 (left-lateral), 0.677 versus 0.652 (right-lateral), and 0.641 versus 0.586 (BIS Quatro): Set C was at or below Set A on every subset (Table 4).

#### Per-subject performance

Decodability varied substantially across held-out participants, and this variation was consistent across montages (Figure 5). Two participants were the poorest decoded in nearly every montage (mean balanced accuracy 0.51 and 0.53 across the five montages), the same two that scored at chance on the full 62-channel array; one never yielded above-chance discrimination on any montage. At the other extreme, three participants were decoded well above chance under every montage (means 0.81–0.96). Reducing the montage rescaled the overall level of performance but did not reorder which participants were decodable.

### 3.4 Variance decomposition of the ketamine effect

We decomposed the per-feature ketamine effect into between-subject, shared, and subject-specific components (Section 2.10), separately for the 5,673 wPLI edges and the 24 band power features.

Across the wPLI edges, the within-subject drug effect exceeded the between-subject variance on 84% of edges (median ratio of between-subject to within-subject variance 0.51); the median edgewise ICC_subject_ was 0.11 across all edges and 0.16 in the alpha band. For the top-20 classifier-weighted alpha edges (Table 7) the median ratio was 0.28 (Figure 6).

**Table 4.** Channel-subset classification performance for the primary random-forest classifier. Balanced accuracy and ROC-AUC are the mean across the ten leave-one-subject-out folds; *p*_bacc_ and *p*_AUC_ are two-sided permutation *p*-values. *n*_feat_ is the number of features per epoch.

| Subset | Set | $n_{\text{feat}}$ | Balanced accuracy | ROC-AUC | $p_{\text{bacc}}$ | $p_{\text{AUC}}$ |
| --- | --- | --- | --- | --- | --- | --- |
| Both lateral | A | 40 | 0.742 | 0.790 | 0.002 | 0.019 |
|  | B | 135 | 0.482 | 0.435 | 0.701 | 0.295 |
|  | C | 175 | 0.701 | 0.793 | 0.008 | 0.003 |
| Left lateral | A | 20 | 0.748 | 0.817 | 0.007 | 0.001 |
|  | B | 30 | 0.488 | 0.493 | 0.818 | 0.919 |
|  | C | 50 | 0.750 | 0.827 | 0.007 | 0.002 |
| Right lateral | A | 20 | 0.677 | 0.767 | 0.017 | 0.042 |
|  | B | 30 | 0.397 | 0.338 | 0.031 | 0.006 |
|  | C | 50 | 0.652 | 0.724 | 0.058 | 0.071 |
| BIS Quatro | A | 16 | 0.641 | 0.742 | 0.056 | 0.003 |
|  | B | 18 | 0.459 | 0.463 | 0.330 | 0.515 |
|  | C | 34 | 0.586 | 0.680 | 0.244 | 0.098 |

Median drug-effect transferability (*τ*) was 0.046 for the wPLI edges, and the across-subject dispersion exceeded the shared component by a factor of ∼ 10 on 95.5% of edges. For the band-matched band power features (theta/alpha/beta), median *τ* was 0.53 (Figure 3A; per-band values in Figure 3B).

A classifier trained to identify subject identity across drug conditions (train awake / test ketamine and vice versa) reached a mean balanced accuracy of 0.43 for band power and 0.39 for wPLI (chance 0.10; Figure 7).

## 4 Discussion

We set out to test whether the subanaesthetic ketamine state can be decoded from single eyes-closed EEG epochs, and how spectral and phase-based connectivity features compare when used for this purpose. The result is a clear dissociation. Log spectral band power discriminated the ketamine state above chance under leave-one-subject-out cross-validation (balanced accuracy 0.707, ROC-AUC 0.803, *p* = 0.011), whereas weighted phase-lag index (wPLI) connectivity computed on the same epochs was statistically indistinguishable from chance (balanced accuracy 0.474, ROC-AUC 0.489, *p* = 0.563), and combining the two feature sets reduced balanced accuracy relative to band power alone (0.671 vs. 0.707). This dissociation held across three classifier families (random forest, support vector machine, and gradient boosting; Table 3), and was not an artefact of the high dimensionality of the connectivity feature space: when wPLI was restricted to small lateral montages of 18–135 edges, with no dimensionality reduction, it remained at chance across every subset tested. In what follows we argue that the wPLI null reflects a property of the subanaesthetic ketamine state rather than a shortcoming of the measure or the analysis, and that the dissociation is an interpretable signature of the kind of brain-state change the drug produces.

**Fig. 3.**
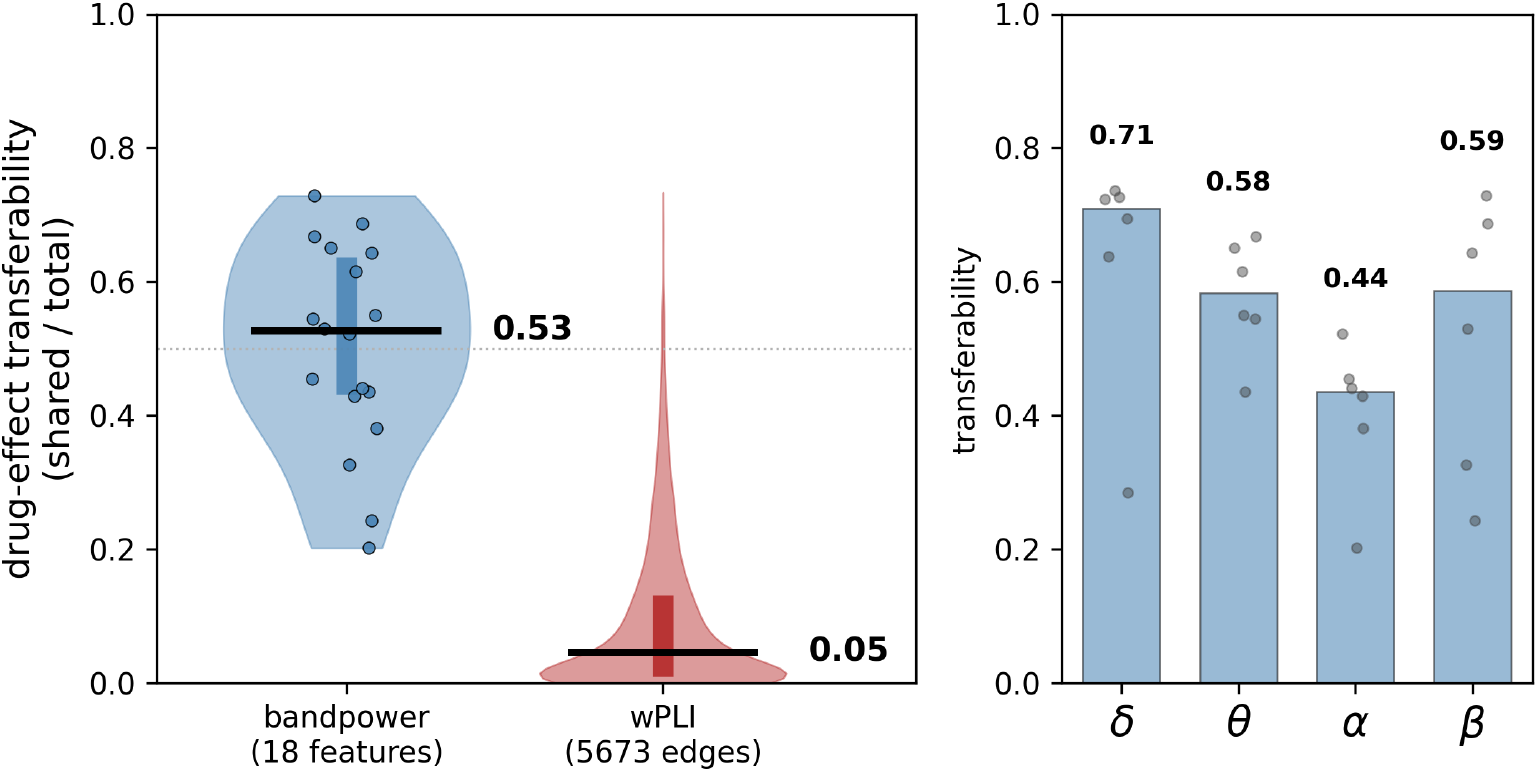
Drug-effect transferability of connectivity versus spectral power, computed per feature. Points show *τ*, the fraction of the awake-versus-ketamine effect shared across subjects (Section 2.10), for all 5,673 wPLI edges and the band-matched band power features (theta/alpha/beta, 18 of 24 features) (left panel), and band power transferability by canonical band (right panel). Thick bars mark medians (0.046 for wPLI, 0.53 for band power in the left panel); the dotted line marks *τ* = 0.5.

### 4.1 Phase-based connectivity indexes the level of consciousness, which subanaesthetic ketamine preserves

The most parsimonious explanation of why band power decodes the ketamine state while wPLI does not begins from what each feature is sensitive to. Subanaesthetic ketamine is defined, in this dataset and in general, by a profound change in the *content* of consciousness, dissociation and mild psychotomimetic effects, with the *level* of consciousness, indexed by responsiveness, left intact (Farnes et al 2020). The original analysis of these recordings makes the same point in complementary terms: spontaneous signal diversity increased under ketamine while the perturbational complexity index, a measure of the brain’s capacity to sustain consciousness, was largely preserved (Farnes et al 2020). Content changed; level did not.

Phase-based functional connectivity appears to track the level rather than the content. In high-density resting EEG during propofol sedation, a different agent acting through GABAergic rather than NMDA mechanisms, Chennu et al showed that alpha-band phase connectivity (measured with the debiased wPLI) tracks behavioural responsiveness. The topological robustness of an individual’s alpha network predicted whether they would lose responsiveness, and connectivity degraded as participants became unresponsive (Chennu et al 2016). Crucially, that study dissociated two axes with two measures; phase-based connectivity tracked responsiveness (the *level* of consciousness), whereas slow–alpha phase-amplitude coupling tracked the blood concentration of the drug (its pharmacological *content* effect). Our design separates the same two axes, but by pharmacology rather than by measure. Spectral power moves with the presence of the drug and is consistent enough across individuals to be decoded, behaving like Chennu et al‘s drug-concentration axis, while undirected phase connectivity, behaving like their responsiveness axis, has little to read out when responsiveness is preserved. On this account the wPLI null is the electrophysiological consequence one would expect from a drug that alters the content of consciousness while leaving its level intact.

This interpretation is reinforced, rather than contradicted, by the anaesthesia literature. The clearest reductions in frontoparietal phase connectivity under ketamine occur at *anaesthetic* doses, when consciousness is lost (Lee et al 2013; Blain-Moraes et al 2014; Vlisides et al 2017), and a comparable breakdown of frontoparietal connectivity occurs under propofol, sevoflurane, and ketamine despite their differing receptor targets (Lee et al 2013). The common factor is therefore the loss of the level of consciousness, not the molecular mechanism. Connectivity collapses when the level drops, regardless of whether the agent is GABAergic or an NMDA antagonist, and is spared when the level is preserved. Subanaesthetic ketamine occupies exactly this latter regime. A secondary, more tentative explanation is that propofol reaches connectivity disruption through graded loss of responsiveness, whereas subanaesthetic ketamine stops short of it; but we regard the level-of-consciousness explanation as primary, precisely because it predicts the anaesthetic-ketamine findings that a purely receptor-based explanation would not.

### 4.2 Why undirected wPLI fails out of sample

The level explanation answers why subanaesthetic ketamine produces little gross, level-tracking network change for a connectivity classifier to exploit. It does not, on its own, explain why the connectivity changes that *are* present fail to support out-of-sample prediction. Our variance decomposition (Section 3.4) provides the specific answer.

The natural explanation was that connectivity is dominated by stable individual differences, a “fingerprint”, that overwhelm a smaller drug effect, a concern grounded in the connectivity-fingerprinting literature (Finn et al 2015; Gratton et al 2018; Mantwill et al 2022). Our data are consistent with the premise of that literature: connectivity is strongly subject-identifying, and subjects could be classified from their wPLI edges well above chance. Between-subject variance, however, did not overwhelm the drug effect; on the contrary, the within-subject ketamine effect exceeded between-subject variance on most edges, and subjects accounted for only a minority of edgewise variance. Identity-dominance is therefore not why the wPLI classifier failed.

The reason is instead the *non-transferability* of the drug effect. Ketamine altered connectivity substantially within each participant, but in a largely subject-specific direction. Only about 5% of the drug effect was shared across subjects, against roughly half for spectral power. Because a subject-aware classifier can exploit only the shared component, an effect this subject-specific leaves almost nothing to generalise, even though it is large within individuals.

This also explains the two auxiliary results. Combining feature classes (Set C) underperformed band power alone on balanced accuracy because concatenating the 5,673 non-generalising wPLI features to the 24 band power features adds dimensions the classifier must learn to down-weight, diluting the spectral signal rather than complementing it. That the combined set retained a ROC-AUC comparable to band power alone, degrading only the threshold-dependent balanced accuracy, is consistent with the wPLI features being uninformative rather than actively misleading. The subject-identification control behaved as it did because both feature classes encode individual identity comparably (balanced accuracy 0.43 for band power, 0.39 for wPLI; Section 3.4), so the asymmetry in *state* decoding reflects how cross-subject-consistent each drug effect is, not how fingerprint-like each feature is.

Two further considerations reinforce this explanation. First, the subanaesthetic connectivity changes that have been reported are not only subtle but largely *directional* : dynamic causal modelling of MEG data shows reduced *effective* frontoparietal coupling (Muthukumaraswamy et al 2015), a directed quantity that an undirected, symmetric measure such as wPLI is not designed to capture, which would further reduce any consistent, transferable signal available to it. Second, the back-projected importance maps indicate that the classifier weighted plausible edges rather than spurious ones. It concentrated on alpha-band edges with a posterior–posterior and posterior–frontal topography (Table 7), the band and locations carrying the largest underlying state difference, yet could not convert that reliance into cross-subject discrimination. The edges it weighted were the right ones; they simply did not change consistently enough across subjects to support generalisation.

This pattern aligns with the most directly comparable EEG study. In low-density EEG during subanaesthetic ketamine, conventional pairwise connectivity was a weak discriminator and a robust, classifiable signal emerged only from higher-order, multivariate interactions (Herzog et al 2024). Our findings extend this to a high-density montage and to a measure, wPLI, chosen specifically for its robustness to volume conduction (Vinck et al 2011), and locate the failure not in spatial sampling but in the subject-specific form of the connectivity drug effect itself. Given the small sample, we advance non-transferability as the best-supported explanation of these data rather than a definitively demonstrated mechanism.

### 4.3 Ruling out mundane explanations

We considered three further mechanisms that might explain the wPLI results, and excluded each. Feature dimensionality is the first: with 5,673 edges and only ∼ 276 epochs, a connectivity classifier might simply be overwhelmed. The channel-subset analysis rules this out, since wPLI remained at chance even with as few as 18 edges and exhaustive hyperparameter tuning (left-lateral 0.488, *p* = 0.818; both-lateral 0.482, *p* = 0.701; BIS-Quatro 0.459, *p* = 0.330). Spatial coverage is the second. The failure was not confined to any one region but held across frontal, temporal, and bilateral montages. One cell fell below chance (right-lateral, 0.397, *p* = 0.031), but this is best read not as an anti predictive connectivity signature but as a small-sample artefact of leave-one-subject-out cross-validation, flagged by the two-tailed permutation test; consistent with this, the cell was below chance for all three classifiers. Third, decodability varied across participants. Two individuals were the least decodable across nearly every montage and feature set, including band power, indicating that they lacked a recoverable epoch-level ketamine signature, spectral or connectivity. Yet the spectral classifier generalised across the remaining majority, whereas wPLI generalised to no one. The dissociation is thus robust to these confounds and localises to the feature class itself.

### 4.4 Implications for clinical monitoring of ketamine states

The spectral signal was not only decodable but spatially sparse, and, critically for any monitoring application, transferable across individuals. A monitor must generalise to patients it was not trained on, and the transferability analysis (Section 3.4) shows that only the spectral signal meets this bar. Ketamine’s effect on connectivity, however large within a given individual, is too subject-specific to support a trained classifier, whereas its effect on band power is largely shared across people. Consistent with this, a five-channel lateral temporal montage recovered the ketamine state as well as the full 62-channel array (left-lateral 0.748/0.817, *p* = 0.007). The discriminative spectral signature is thus spatially sparse: it is carried by temporal-chain electrodes alone, without central, parietal, or midline coverage, and adding the second hemisphere did not improve balanced accuracy (both-lateral 0.742), so a single temporal chain already carries most of the available signal.

This bears on the monitoring of the subanaesthetic ketamine state, an active concern as the drug becomes a mainstream treatment for depression (Chan et al 2025). Unlike depth-of-anaesthesia indices such as the BIS, which are calibrated for the loss of consciousness and are widely reported to be unreliable indices of the ketamine state (Hans et al 2005; Roffey et al 2000), monitoring the conscious, subanaesthetic state is a distinct problem, and our results speak to it directly. The footprint of a forehead sensor strip retains decodable information about the ketamine state, but only through spectral features rather than a proprietary index. A four-channel frontal montage approximating the BIS sensor strip yielded a significant probabilistic discriminator (ROC-AUC 0.742, *p* = 0.003) but not a significant binary classifier at the default threshold (balanced accuracy 0.641, *p* = 0.056). This split between metrics is itself informative: ROC-AUC integrates over all decision thresholds, whereas balanced accuracy is evaluated at the fixed 0.5 threshold and is more sensitive to class imbalance and small-sample miscalibration, so the frontal montage retains usable but reduced spectral information about the ketamine state. Five temporal-chain sensors, a comparable footprint, carried substantially more decodable signal. The practical message is therefore that electrode *placement* matters more than channel count, that spectral power is the workhorse feature, and that phase-based connectivity adds nothing usable in this setting. This aligns with the independent observation that resting subanaesthetic ketamine, without accompanying transitions in the level of consciousness, offers limited state-related variability for connectivity or complexity measures to exploit (Chan et al 2025).

### 4.5 Limitations

Several limitations bound these conclusions. The sample is small (*N* = 10) and inherited rather than designed, and the design is open-label and within-subject; our claims concern the decodability of the state, not the antidepressant or subjective effects of the drug. The spectral signature we exploited reproduced, at canonical-band resolution and on Laplacian-derived CSD estimates, the broadband power reductions previously reported for this dataset (Farnes et al 2020). Absolute power was nonetheless highly variable across subjects, particularly in the beta range, consistent with beta-band power being both strongly trait-like and heritable (Smit et al 2005) and the most susceptible of the lower bands to arousal- and electromyography-related contamination (Goncharova et al 2003); this dispersion is independent of the within-subject drug effect the classifier exploits. The canonical subanaesthetic signature also includes a gamma-range power increase (Muthukumaraswamy et al 2015; de la Salle et al 2016), which we did not recover. No significant high-frequency cluster emerged (cluster *p* = 0.33), in keeping with the known difficulty of recovering scalp gamma and our conservative cluster threshold. We therefore make no claim about gamma in this dataset. Most importantly, our central claim is deliberately narrow. It is that *undirected* phase-based connectivity does not support *out-of-sample, single-epoch* decoding of the *subanaesthetic* ketamine state in this dataset. It is not a claim that ketamine leaves functional connectivity unchanged. Group-level and directional analyses do detect subanaesthetic connectivity effects (Muthukumaraswamy et al 2015), fMRI connectivity classifies ketamine from saline (Joules et al 2015), though fMRI is sensitive to subcortical and deep-network contributions that scalp-EEG wPLI cannot resolve, so a positive fMRI result does not imply the same signal is accessible to surface EEG, and connectivity collapses under anaesthetic-dose ketamine (Lee et al 2013); our result locates the limits of *undirected, predictive, subject-generalising* connectivity specifically, and the gap between this and the positive group-level literature is itself the finding. The propofol comparison we draw is a conceptual convergence across datasets and drugs, not a controlled within-study contrast, and would be strengthened by a design in which dose is titrated across the threshold of responsiveness within the same NMDA-antagonist agent.

## Statements and Declarations

### Ethics approval

This study is a secondary analysis of a publicly available, de-identified dataset. The original data collection was approved by Norwegian Regional Committees for Medical and Health Research Ethics (reference 2015/1520/REK Sør-Øst A) and was performed in accordance with the ethical standards laid down in the 1964 Declaration of Helsinki and its later amendments; full details are reported in Farnes et al (2020). No new data were collected for the present study, and no additional ethics approval was required for this reanalysis.

### Consent to participate

The present study analysed only de-identified data and involved no new contact with participants.

### Consent to publish

The present study reports no identifying information about individual participants.

### Data availability

The EEG data analysed in the present study were collected and released by Farnes et al (2020) and are publicly available from the Dryad Digital Repository (10.5061/dryad.j9kd51c9q). No new data were generated in the course of this study.

### Code availability

All analysis code required to reproduce the results reported here is publicly available on GitHub.

### Author contributions

All authors contributed to the study conception and design. HS performed the software implementation, formal analysis, data curation, and visualization, and wrote the first draft of the manuscript. FVW supervised the project. Both authors critically reviewed and edited the manuscript, and read and approved the final version.

## Appendix A Dataset composition

**Table 5.** Per-recording epoch counts in the analysed dataset. All 276 epochs released by Farnes et al passed the 250 µV peak-to-peak threshold applied in the present study and were retained.

| Subject | Condition | Retained epochs |
| --- | --- | --- |
| sub-01 | awake | 14 |
| sub-01 | ketamine | 11 |
| sub-02 | awake | 14 |
| sub-02 | ketamine | 16 |
| sub-03 | awake | 15 |
| sub-03 | ketamine | 17 |
| sub-04 | awake | 14 |
| sub-04 | ketamine | 13 |
| sub-05 | awake | 13 |
| sub-05 | ketamine | 11 |
| sub-06 | awake | 14 |
| sub-06 | ketamine | 14 |
| sub-07 | awake | 14 |
| sub-07 | ketamine | 11 |
| sub-08 | awake | 14 |
| sub-08 | ketamine | 14 |
| sub-09 | awake | 15 |
| sub-09 | ketamine | 14 |
| sub-10 | awake | 14 |
| sub-10 | ketamine | 14 |
| **Total** |  | **276** |

## Appendix B Per-subject spectral and decoding results

**Fig. 4.**
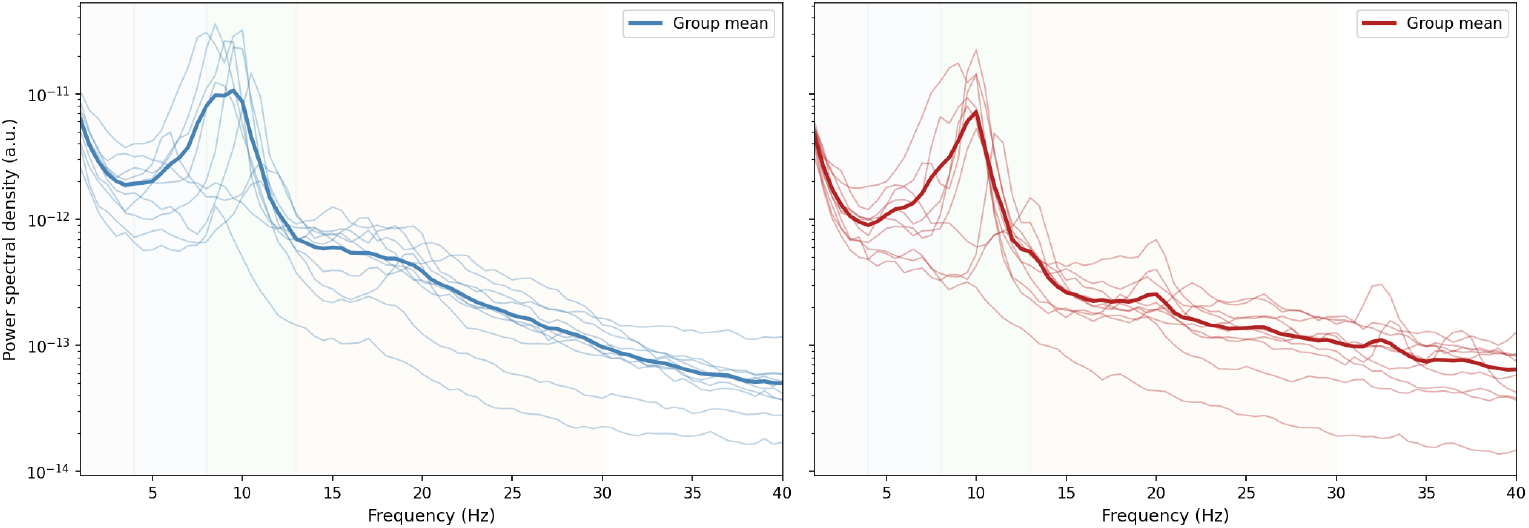
Per-subject power spectral densities for the eyes-closed recordings, separately for the awake (left, blue) and ketamine (right, red) conditions. Thin lines: individual participants (*N* = 10); thick lines: across-subject mean. Background shading marks canonical frequency bands. Power is in arbitrary units (a.u.) because the data are surface-Laplacian-derived current-source-density estimates. Every participant exhibited the ketamine-induced suppression in the theta and beta ranges identified at the group level (Figure 1, Table 1); inter-subject variability in absolute power was substantial, particularly in the beta range

**Fig. 5.**
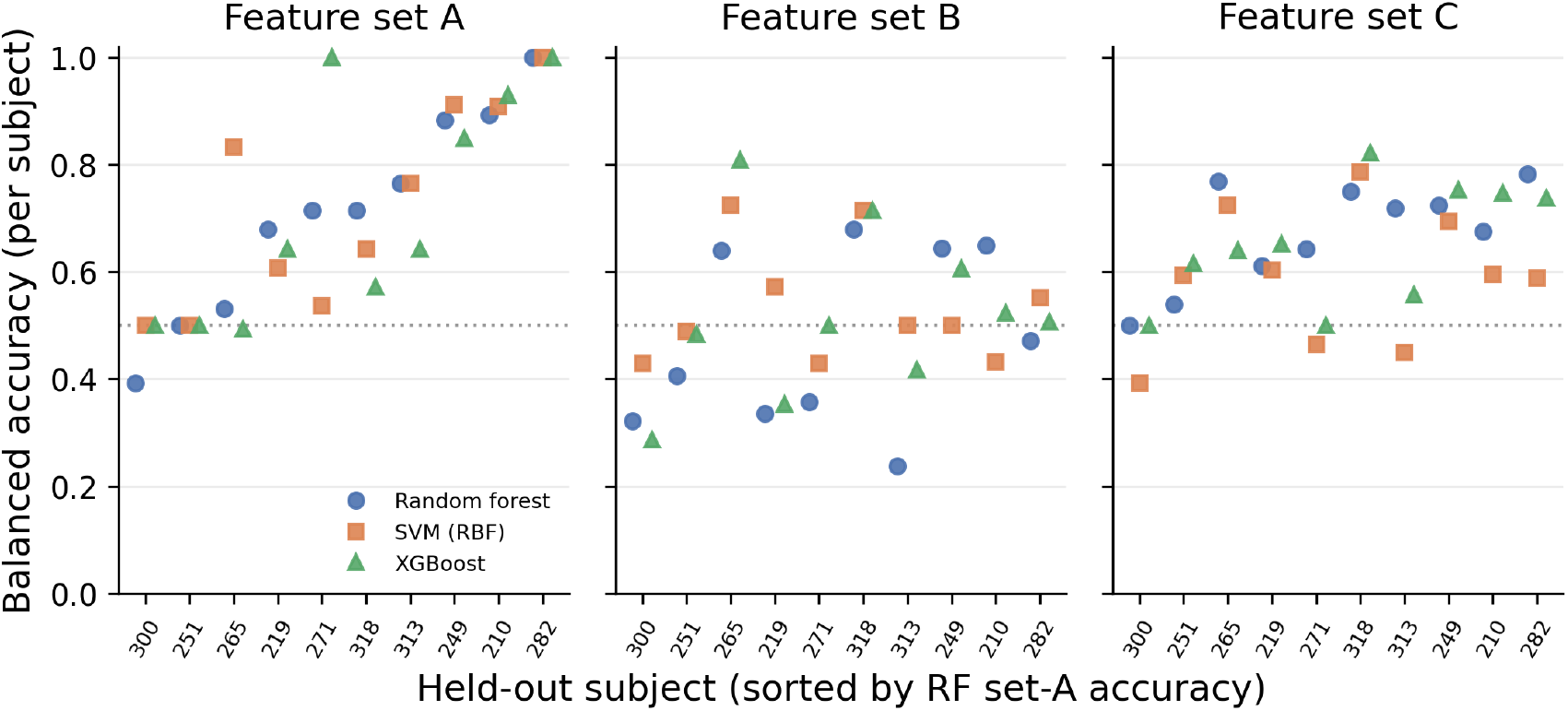
Per-subject balanced accuracy under leave-one-subject-out cross-validation (full 62-channel montage). Because each outer fold holds out exactly one subject, every fold is a single-subject test, and each marker gives one held-out subject’s balanced accuracy. Panels correspond to the three feature sets (A, band power; B, wPLI; C, A+B). Within a panel the x-axis lists the ten held-out subjects, ordered by ascending random-forest set-A accuracy so that the poorest-decoded subjects appear on the left; the same subject order is used in all three panels. Marker shape and colour denote the classifier (random forest, circles; RBF-SVM, squares; XGBoost, triangles). The dotted line at 0.5 indicates chance. For band power (A) all three classifiers decode the drug state well above chance in the majority of subjects, but fail jointly on subjects 300 and 251 (balanced accuracy ≤ 0.5; sensitivity 0, i.e. no ketamine epoch is flagged), indicating that these failures are subject-specific rather than an artefact of a particular algorithm. Connectivity (B) sits near chance for essentially every subject and classifier, whereas the combined set (C) tracks band power. The full-montage results summarised here are discussed in Section 3.2

## Appendix C Subject-identity controls

**Fig. 6.**
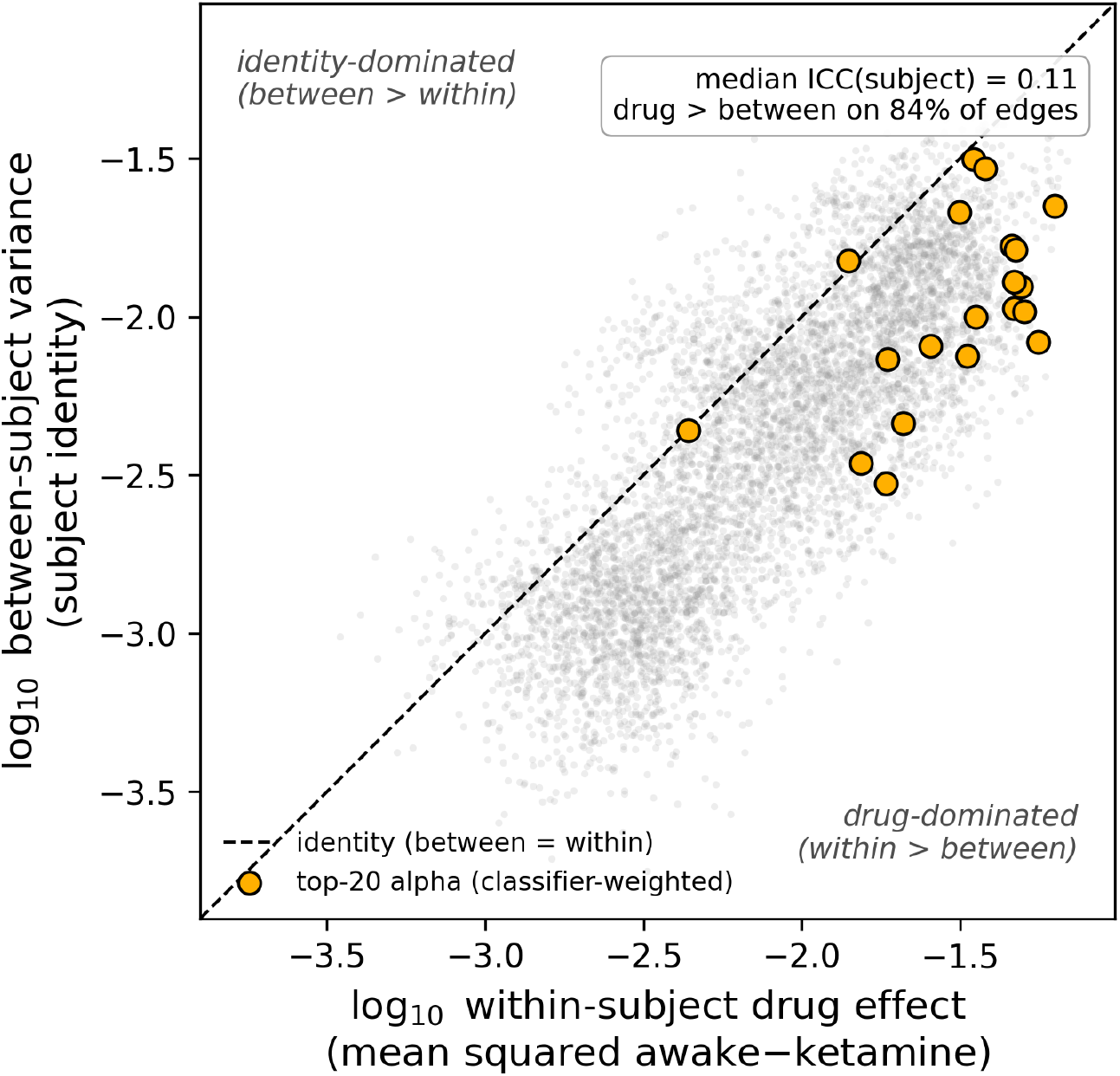
Between-versus within-subject variance of wPLI edges. Each grey point is one of the 5,673 edges; axes show the within-subject drug effect (mean squared awake−ketamine difference) against the between-subject variance (subject identity), both on a log_10_ scale. The dashed line marks equality. Points below the line are drug-dominated (the within-subject ketamine effect exceeds between-subject variance), which holds for 84% of edges; the median edgewise ICC_subject_ is 0.11. Amber points mark the top-20 classifier-weighted alpha edges (Table 7), which are if anything more drug-dominated than average—identity does not dominate the connectivity feature space

**Fig. 7.**
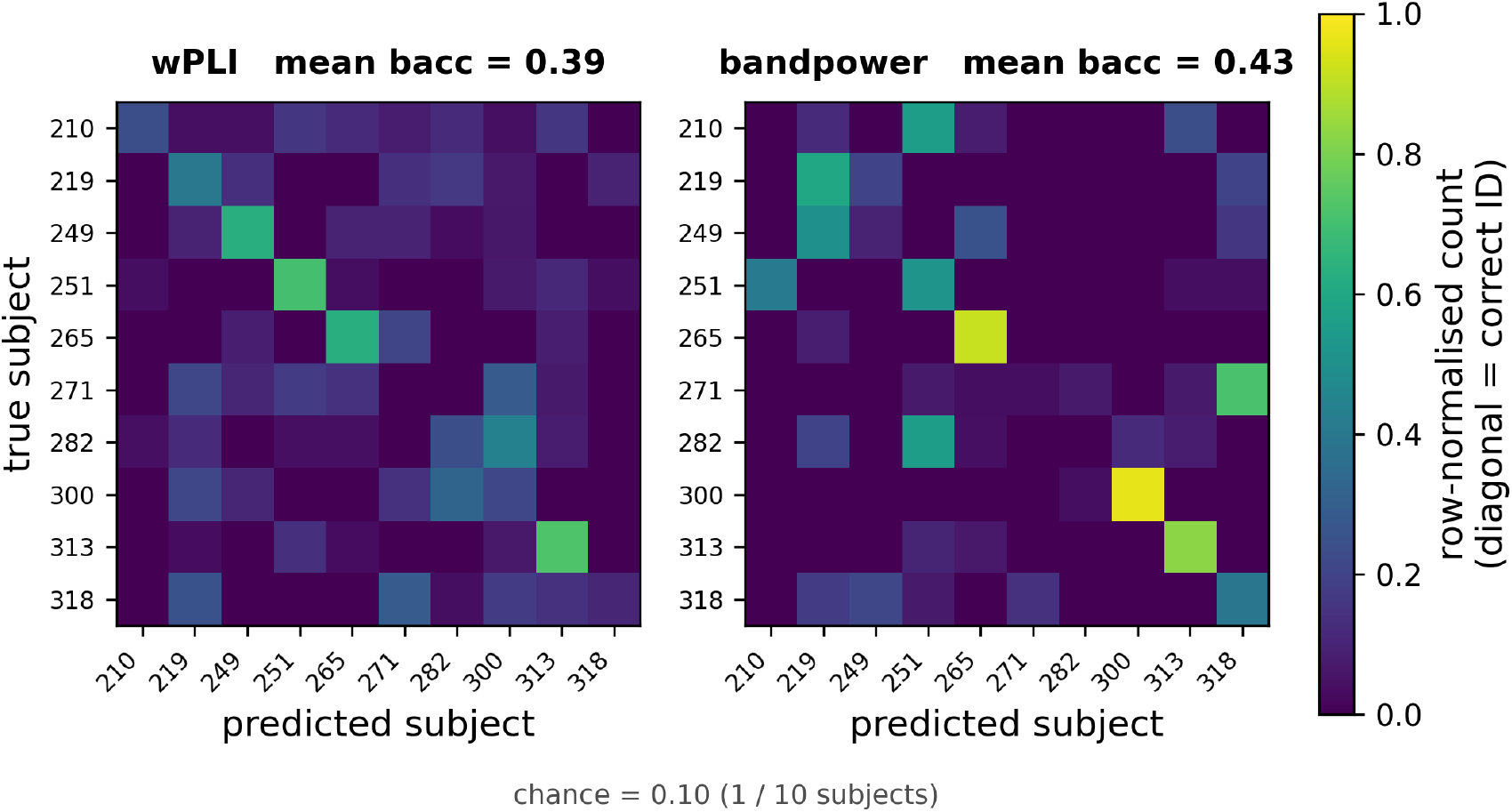
Subject-identification control. Row-normalised confusion matrices for a random-forest classifier trained to predict subject identity (10 classes) from wPLI edges (left) and band power features (right), evaluated across drug conditions (train awake / test ketamine and vice versa) so that identity decoding cannot exploit within-recording leakage. Both feature classes identify subjects well above the 0.10 chance level (mean balanced accuracy 0.39 for wPLI, 0.43 for band power); band power is if anything the more subject-identifying of the two. Differential identity-dominance therefore cannot explain why band power decodes the drug state while wPLI does not

## Appendix D Back-projected wPLI edge importance

We report the spatial distribution of random-forest impurity-based importances for the full-62-channel wPLI classifier (Section 3.2), projected back into the original 5,673-edge space using the squared-loadings method described in Section 2.9. Because the underlying classifier did not exceed chance (*p*_perm_ = 0.563 on the two-sided permutation test), the patterns reported here cannot be interpreted as evidence of class-discriminative connectivity in the present data. They are provided to document what the classifier weighted and to support potential follow-up analyses in larger samples.

Figure 8 shows the back-projected importance, averaged across the ten outer folds, as a 62 × 62 matrix for each of the three frequency bands. Theta and alpha bands showed broadly distributed importance across most electrode pairs; the beta band was sparser, with overall lower mean magnitudes. When edges were ranked by mean importance, however, the top of the distribution was dominated by the alpha band: all 20 of the highest-ranked edges across the three bands belonged to alpha (Table 7). Spatially, these edges clustered into two overlapping motifs: dense posterior–posterior coupling between occipital and parieto-occipital electrodes (e.g. Oz–PO7, O1–PO7, PO4–POz, Oz–O1), and long-range posterior–frontal edges (e.g. P8–FC3, POz–Fp2, P1–FT8). This is consistent with alpha being the dominant resting rhythm in eyes-closed EEG and with the alpha-band suppression effect identified in Section 3.1: the classifier weighted the band and spatial topography where the largest underlying state difference was located, even though it was unable to translate this attention into reliable cross-subject discrimination. The failure to generalise may reflect that the relevant alpha-band coupling differences are too small relative to between-subject variability, or that wPLI on Laplacian-derived CSD estimates is suboptimally sensitive to whatever phase relationship actually changes under ketamine. Both possibilities warrant investigation in larger samples.

**Fig. 8.**
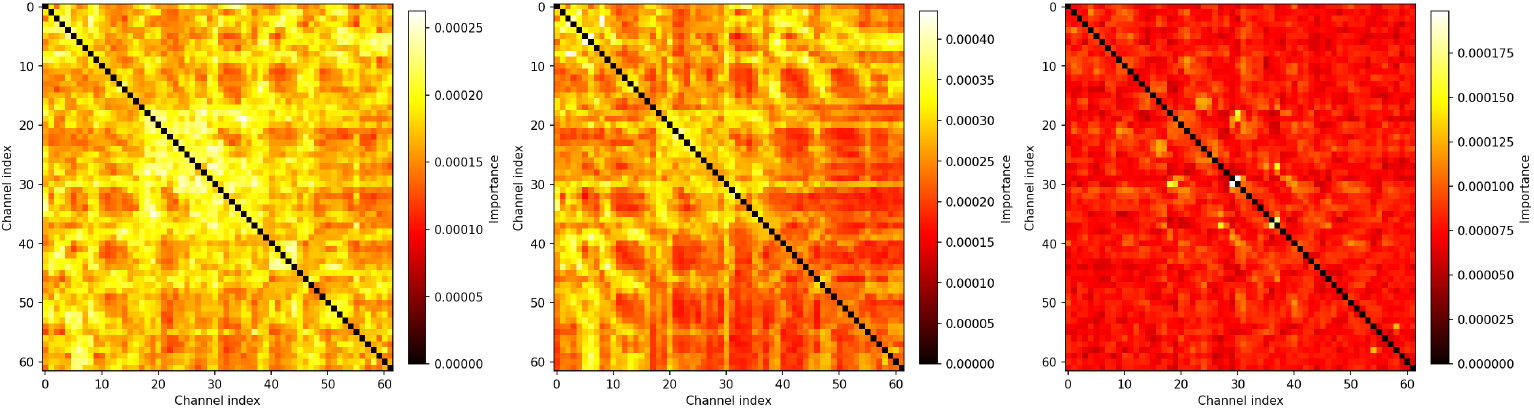
Back-projected random-forest impurity importance for the full-62-channel wPLI classifier, in the original 5,673-edge space. Each panel shows a 62 *×* 62 symmetric matrix for one frequency band; the diagonal is masked. Importance values are averages across the ten outer cross-validation folds, with squared loadings used to avoid sign cancellation. Higher values indicate edges to which the classifier assigned more importance. Channel indices 0–61 are ordered approximately posterior (Iz, index 0) to anterior (Fp1, index 61); the full index-to-electrode mapping is given in Table 6. As the underlying classifier did not perform above chance (Section 3.2), these maps are presented for methodological completeness only

**Table 6.** Channel index → 10–10 electrode name for the 62 *×* 62 wPLI edge-importance heatmaps (Figure 8, Feature set B). Indices 0–61 are the native channel order of the source Dryad recordings. Edge labels in Table 7 follow the same ordering.

| Idx | Ch. | Idx | Ch. | Idx | Ch. | Idx | Ch. |
| --- | --- | --- | --- | --- | --- | --- | --- |
| 0 | Iz | 16 | P5 | 32 | C2 | 48 | F6 |
| 1 | O2 | 17 | P7 | 33 | Cz | 49 | F4 |
| 2 | Oz | 18 | TP10 | 34 | C1 | 50 | F2 |
| 3 | O1 | 19 | TP8 | 35 | C3 | 51 | Fz |
| 4 | PO8 | 20 | CP6 | 36 | C5 | 52 | F1 |
| 5 | PO4 | 21 | CP4 | 37 | T7 | 53 | F3 |
| 6 | POz | 22 | CP2 | 38 | FT8 | 54 | F5 |
| 7 | PO3 | 23 | CPz | 39 | FC6 | 55 | F7 |
| 8 | PO7 | 24 | CP1 | 40 | FC4 | 56 | AF4 |
| 9 | P8 | 25 | CP3 | 41 | FC2 | 57 | AF7 |
| 10 | P6 | 26 | CP5 | 42 | AF8 | 58 | AF3 |
| 11 | P4 | 27 | TP7 | 43 | FC1 | 59 | Fp2 |
| 12 | P2 | 28 | TP9 | 44 | FC3 | 60 | Fpz |
| 13 | Pz | 29 | T8 | 45 | FC5 | 61 | Fp1 |
| 14 | P1 | 30 | C6 | 46 | FT7 |  |  |
| 15 | P3 | 31 | C4 | 47 | F8 |  |  |

**Table 7.** Top 20 wPLI edges ranked by mean back-projected random-forest impurity importance across the ten outer cross-validation folds, for the full-62-channel wPLI classifier (Set B, Section 3.2). Edges are listed in descending order of importance. Channel labels follow the international 10–10 system. As the underlying classifier did not perform above chance (balanced accuracy 0.474, *p*_perm_ = 0.563; Section 3.2), this ranking should be treated as hypothesis-generating only; no claim is made that these edges carry class-discriminative information in the present data.

| Rank | Band | Channel pair | Mean importance |
| --- | --- | --- | --- |
| 1 | Alpha | Oz–PO7 | $4.35 \times 10^{-4}$ |
| 2 | Alpha | PO4–POz | $4.24 \times 10^{-4}$ |
| 3 | Alpha | O1–PO7 | $4.15 \times 10^{-4}$ |
| 4 | Alpha | Oz–O1 | $4.02 \times 10^{-4}$ |
| 5 | Alpha | P8–FC3 | $3.97 \times 10^{-4}$ |
| 6 | Alpha | POz–FC6 | $3.95 \times 10^{-4}$ |
| 7 | Alpha | CP4–CP2 | $3.91 \times 10^{-4}$ |
| 8 | Alpha | O2–PO8 | $3.87 \times 10^{-4}$ |
| 9 | Alpha | PO8–FC3 | $3.82 \times 10^{-4}$ |
| 10 | Alpha | POz–Fp2 | $3.81 \times 10^{-4}$ |
| 11 | Alpha | O1–FC4 | $3.78 \times 10^{-4}$ |
| 12 | Alpha | O2–FC3 | $3.76 \times 10^{-4}$ |
| 13 | Alpha | Iz–Cz | $3.75 \times 10^{-4}$ |
| 14 | Alpha | P1–FT8 | $3.73 \times 10^{-4}$ |
| 15 | Alpha | Iz–C1 | $3.73 \times 10^{-4}$ |
| 16 | Alpha | P8–C3 | $3.68 \times 10^{-4}$ |
| 17 | Alpha | POz–P6 | $3.65 \times 10^{-4}$ |
| 18 | Alpha | PO4–P6 | $3.64 \times 10^{-4}$ |
| 19 | Alpha | P8–C6 | $3.64 \times 10^{-4}$ |
| 20 | Alpha | P3–FC6 | $3.63 \times 10^{-4}$ |

